# Anticipatory neural activity improves the decoding accuracy for dynamic head-direction signals

**DOI:** 10.1101/439950

**Authors:** Johannes Zirkelbach, Martin Stemmler, Andreas V.M. Herz

## Abstract

Insects and vertebrates harbor specific neurons that encode the animal’s head direction (HD) and provide an internal compass for spatial navigation. Each HD cell fires most strongly in one preferred direction. As the animal turns its head, however, HD cells in rat anterodorsal thalamic nucleus (ADN) and other brain areas fire already before their preferred direction is reached, as if the neurons anticipated the future head direction. This phenomenon has been explained at a mechanistic level but a functional interpretation is still missing. To close this gap, we use a computational approach based on the animal’s movement statistics and a simple model for the behavior of the ADN head-direction network. Network activity is read out using population vectors in a biologically plausible manner, so that only past spikes are taken into account. We find that anticipatory firing improves the representation of the *present* HD by reducing the motion-induced temporal bias inherent in causal decoding. The amount of anticipation observed in ADN enhances the precision of the HD compass read-out by up to 40%. In addition, our framework predicts that neural integration times not only reflect biophysical constraints, but also the statistics of natural stimuli; anticipatory tuning should be found whenever neurons encode sensory signals that change gradually in time.

**Significance statement:** Across different brain regions, populations of noisy neurons encode dynamically changing stimuli. Decoding a time-varying stimulus from the population response involves a trade-off: For short read-out times, stimulus estimates are unreliable as the number of stochastic spikes will be small; for long read-out times, estimates are biased because they lag behind the true stimulus. We show that optimal decoding relies not only on finding the right read-out time window, but requires neurons to anticipate future stimulus values. We apply this framework to the rodent head-direction system and show that the experimentally observed anticipation of future head directions can be explained at a quantitative level from the neuronal tuning properties, the network size, and the animal’s head-movement statistics.

## Introduction

Spatial orientation requires information about one’s current position and heading direction. In rodents, positional information is provided by place cells (O’Keefe and Dostrovsky, 1971), grid cells (Hafting et al., 2005), boundary vector cells (Barry et al., 2006) and further spatially modulated neurons. As in other vertebrate (Finkelstein et al., 2015) and insect (Seelig and Jayaraman, 2015) species, directional information is represented by head direction (HD) cells, which reside in various brain areas and are particular common along the Papez circuit (Taube, 2007).

HD cells integrate motor efference copies with vestibular, visual and proprioceptive signals, and are maximally active when the animal’s head points into a cell-specific preferred direction (Taube, 2007). These directions are uniformly distributed and rotate in close synchrony with rotated salient visual cues. During navigation in darkness, direction-specific firing is maintained but the preferred directions drift over time (Taube et al., 1990b). Optogenetically interfering with the HD system impairs homing, demonstrating that the HD system plays a crucial role for spatial navigation (Butler et al., 2017).

Conceptually, one can arrange the preferred directions of HD cells on a ring that is anchored to the local environment (Figure 1). When the animal turns its head, the packet of active neurons is updated such that those cells are most active whose preferred directions match the animal’s new HD. As the HD can be read out from the position of this activity packet along the ring, the HD system is often referred to as an internal or neural compass. Unlike the needle of a physical compass, which points towards magnetic north, the preferred directions of HD cells are determined by local cues and vary from environment to environment.

**Figure 1.**
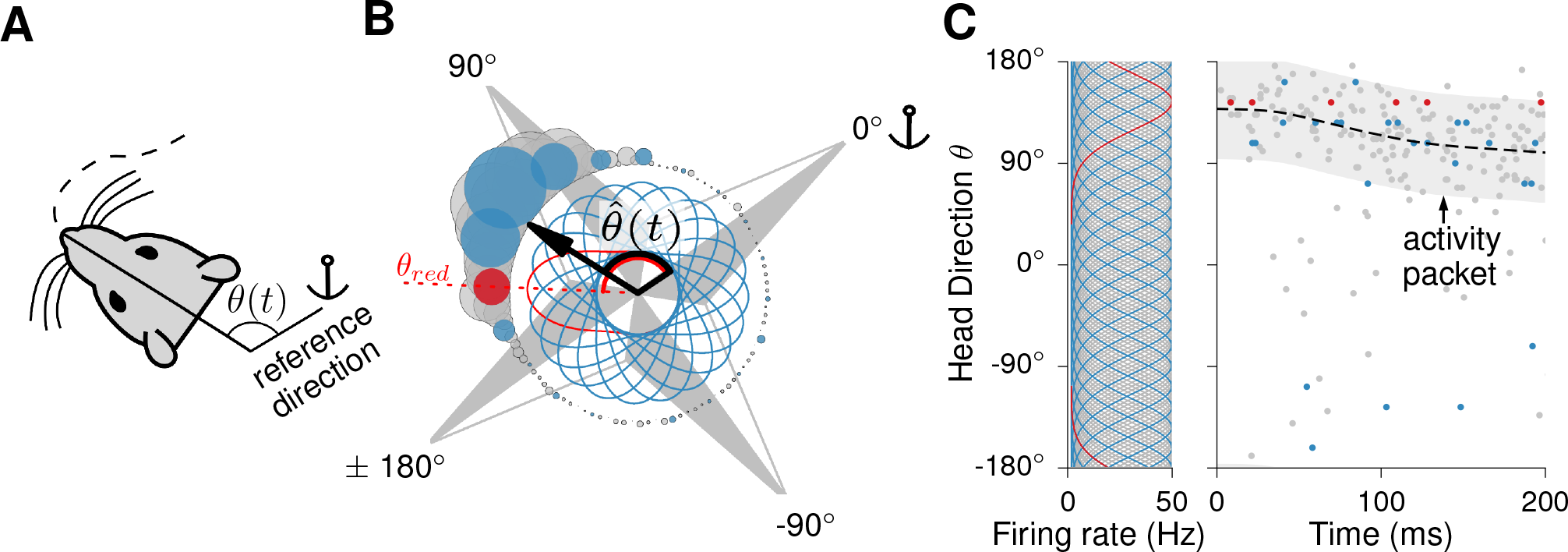
Population encoding and decoding of head direction. **(A)** The head direction (HD) system acts as an internal compass, encoding the time-dependent azimuth *θ*(*t*) of the animal’s head with respect to a landmark-anchored reference direction. **(B)** Polar plot of idealized HD tuning curves and activities. HD cells are broadly tuned (∼ 90°) so that about 25% of all cells are active at any moment. The tuning curves – highlighted for twenty neurons by the inner blue and red lines – have an elevated baseline (innermost circle) and their maximum is achieved at the cell’s preferred direction, as illustrated for the red cell. The activation levels of the different cells at a particular time *t* are denoted by the disk radii. The population vector 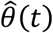 is calculated from these stochastic activities and provides the time-dependent direction estimate. **(C)** Temporal evolution of the HD activity packet for a small clockwise head movement *θ*(*t*), shown as dashed line. Blue, red and grey dots in the raster plot represent spikes generated with inhomogeneous Poisson processes whose rates are determined by the current head direction via the tuning curves shown on the left. Model parameters: Peak firing rate: 50 Hz, background firing rate: 2 Hz.

How accurately can an animal determine its head direction from this compass? Typical tuning widths of individual HD cells (Figure 1B, Figure 2) range from 30° to 70° (Table 1) – too large for accurate navigation on the basis of single-cell responses. However, combining the information provided by all simultaneously active HD cells leads to a far more reliable estimate. To understand HD coding at the network-level, we construct a computational model, read out its activity using population-vector decoding (Georgopoulos et al., 1983; Seung and Sompolinsky, 1993; Salinas and Abbott, 1994), and study the accuracy and bias of this neural compass. For concreteness, we focus on the anterodorsal thalamic nucleus (ADN), which is a core region of the HD system (Taube, 2007) and contains the highest fraction of HD cells (Taube, 2003).

**Figure 2.**
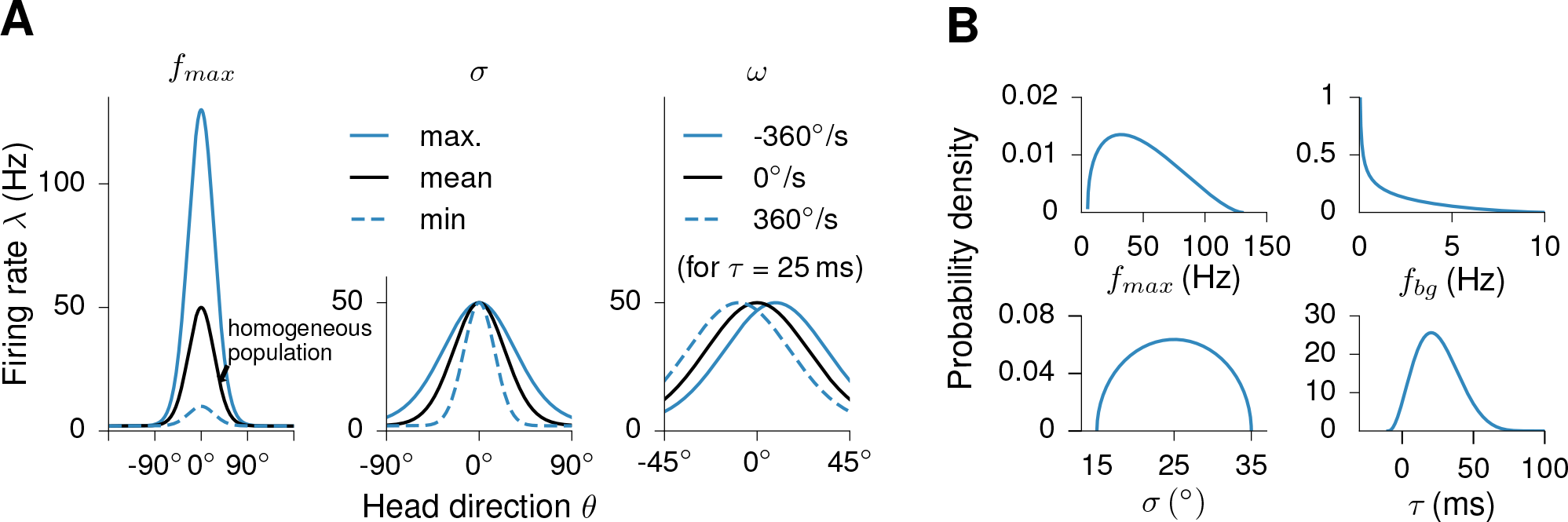
Tuning curves: shapes and parameter ranges. **(A)** Left and middle panel: Illustration of von Mises tuning curves for different choices of the minimum, mean, and maximum values of the peak firing rate *f*_*max*_ and tuning width *σ* (for anticipatory time delay *τ* = 0). Right panel: Shift of a tuning curve during fast head turns (angular head velocity: ±360°/s); note the different scale on the *θ*-axis. **(B)** Parameter distributions from which the tuning curves were sampled to generate inhomogeneous populations (beta distributions whose ranges were adapted to the experimental observations). The distributions have means, standard deviations, and range values as shown in Table 1.

**Table 1.** Properties of head direction cells in the anterodorsal thalamic nucleus. Summary of HD cell parameters as reported in the ADN literature (Blair and Sharp, 1995; Goodrich and Taube, 1997; Blair et al., 1997, 1998; Stackman and Taube, 1998; Taube and Muller, 1998; Taube, 2007; Clark et al., 2009, 2012). *f*_*max*_ and *f*_*bg*_ denote the peak and background firing rates, respectively, *σ* stands for the tuning width, and *τ* is the anticipatory time interval. The concentration parameter *κ* of von Mises tuning curves is given by *κ* = (*σ·π*/180°)^−2^ so that *κ* ranges from about 2.7 (for *σ* = 35°) to about 14.6 (for *σ* = 15°) and the mean *σ* value corresponds to*κ* ≈ 5.25.

| Parameter | Mean value | Standard deviation | Range |
| --- | --- | --- | --- |
| $f_{max}$ | 50 Hz | $\pm 27$ Hz | 5 - 130 Hz |
| $f_{bg}$ | 2 Hz | $\pm 2$ Hz | 0 - 10 Hz |
| $\sigma$ | $25^\circ$ | $\pm 5^\circ$ | $15^\circ - 35^\circ$ |
| $\tau$ | 25 ms | $\pm 15$ ms | -10 - 100 ms |

During foraging, the animal’s head direction becomes a time-varying quantity. In this case, causal decoding – for which only spikes from the past are used – biases the population vector, as we will show. To counteract this temporal bias, whose size is determined by the time course of the HD trajectory, the decoder can reduce the read-out time, yet shorter read-out times result in higher variability.

Interestingly, HD cells in the ADN fire in an anticipatory manner: during head turns, their preferred directions shift such that the cells fire earlier than without movement – as if they encoded the future HD (Blair and Sharp, 1995). Our decoding approach suggests an alternative interpretation: anticipatory firing improves the representation of the *present* HD by compensating the motion-induced bias inherent in the population vector. If cells anticipate the HD by 25 ms, as observed in the ADN, the HD compass read-out improves by up to 40%. Moreover, across the layers within the HD circuit, the anticipatory time intervals should be staggered to optimize the population-vector codes, which is consistent with experimental observations (see, e.g., Taube, 2007). Similar phenomena are to be expected in other multi-layered neural systems that encode stimuli that vary smoothly in time.

## Materials and Methods

### Head direction trajectories

The head direction trajectories *θ*(*t*) were recorded in the Buzsáki lab (Mizuseki et al., 2009) and are publicly available at http://dx.doi.org/10.6080/K0Z60KZ9. The tracking data (file extension *.whl) are from rats foraging on square platforms (side length: 120 or 180 cm). The files contain time traces (sampling rate: 39.6 Hz) of the x- and y-coordinates of two LEDs mounted on the rats’ head stage. To facilitate analyses at a fine temporal scale, we linearly interpolated the line segments between the data points to obtain an artificial sampling rate of 1 kHz. The time course of the angle between the vector connecting the LEDs (caudal to rostral) and the positive x-axis defines the HD trajectory *θ*(*t*) with −*π* ≤ *θ*(*t*) < *π*. To avoid erroneous data points, we cut each tracking file at time points where tracking data were missing and only kept trajectory segments during which the distance between the LEDs was between 5 cm and 16 cm. Lower distances result from strong head tilts (e.g., during eating or rearing), larger distances indicate recording errors (distances cannot be larger than the distance between the LEDs). From the remaining segments only those longer than 10 s were kept for further analysis. Two of those segments were sorted out manually because of recording irregularities. In sum, this procedure resulted in 1163 segments. A representative segment with a duration *T*_*E*_ of about 90 s (see Figure 3) was chosen for the Monte Carlo simulations described below. This segment length is a compromise between sampling enough of the (foraging) behavior of the animal while keeping the computation time at a moderate level, and leads to representative statistics as shown in Figure 7. Note that we will mainly use the radian measure for angles in the methods section. In the figures and the results section, we will state the angles in the degree measure for a more intuitive reading.

**Figure 3.**
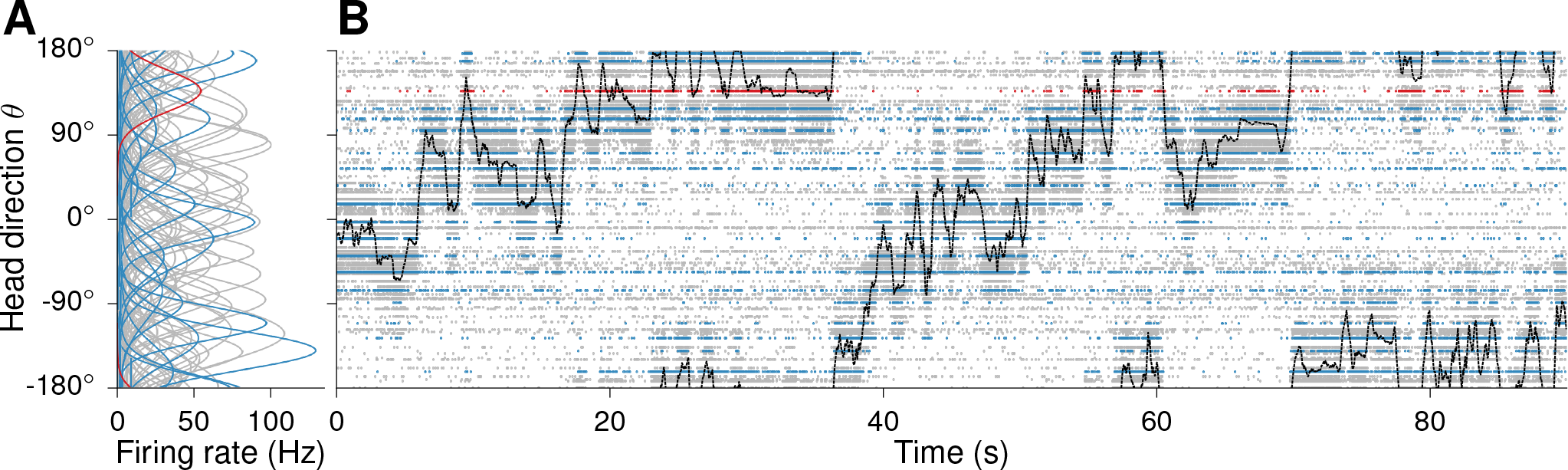
Encoding model and head direction trajectory. **(A)** Tuning curves, modeled as von Mises functions, of a realistic inhomogeneous population of head direction (HD) cells in the anterodorsal thalamic nucleus (N = 100). The parameters of the von Mises functions were drawn from the distributions shown in Figure 2. **(B)** Time-dependent head direction *θ*(*t*) obtained from a two-LED recording of a foraging rat (black line). In addition, spike times from one realization of inhomogeneous Poisson processes based on the rat’s movement trajectory are shown for each neuron.

**Figure 4.**
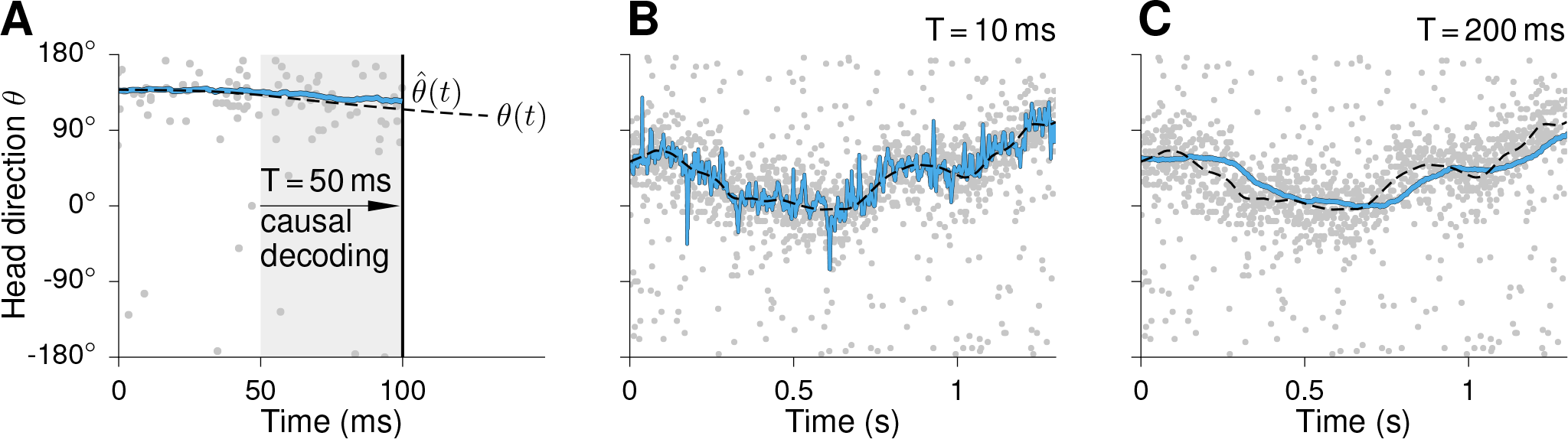
Causal decoding and effect of the spike-count window. **(A)** Within a causal framework, only spikes from the past can be used to estimate the encoded variable. To decode the time-dependent head direction *θ*(*t*) shown as dashed black line at the time of the solid black line, one might, for example, rely on spikes generated within the gray area. The blue line depicts the time-dependent estimate 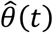 which was calculated from the shown spike responses using the population vector decoder. **(B)** and **(C)** Estimated head directions 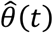 using a short (B) and a long (B) spike-count window *T*. For the short spike-count window 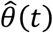 jitters strongly around the true value *θ*(*t*), whereas for longer *T* the estimated head direction is much smoother but lags behind the true trajectory because spikes from the more distant past are used.

To analyze the effect of anticipatory firing, the time-dependent angular head velocity (AHV) *ω*(*t*) needs to be known. We numerically calculated *ω*(*t*) from the experimentally obtained HD trajectories *θ*(*t*) by central differences i.e., *ω*(*t*_*i*_) = [*θ*(*t*_*i*+1_) − *θ*(*t*_*i*−1_)]/[*t*_*i*+1_ − *t*_*i*−1_], to make sure that position and velocity samples are synchronized. Note that in this case the circular trajectory *θ*(*t*) was unfolded at the wrap-around *θ*(*t*) = ±*π*.

### Spike train model

Each model neuron *j* generates an inhomogeneous Poisson spike train with time-dependent rate *λ*_*j*_(θ(*t*)). Accordingly, the number of spikes *k*_*j,T*_(*t*) fired in a time window of length *T* ending at *t* is given by a Poisson distribution with time-dependent mean 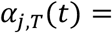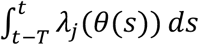,

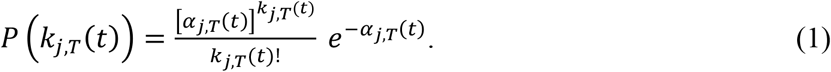

The population response in that time window is given by the spike-count vector 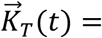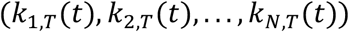. Assuming independence, the probability 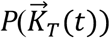 to observe a specific spike-count vector factorizes, i.e., 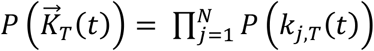. Figure 3 shows a raster plot of HD spike trains generated by this process.

### Homogeneous and inhomogeneous cell populations

Our mathematical analysis focuses on homogeneous populations of HD cells. In these model populations, neurons have identical von Mises tuning curves *λ*_*i*_(*θ*), given by Equation (17) in the Results section, which differ only by their equidistant preferred directions *θ*_*i*_. All other parameters (maximum firing rate *f*_*max*_, background firing rate *f*_*bg*_, inverse tuning width *κ* and anticipatory time interval *τ*) are chosen as the mean values from Table 1 which contains average experimental HD tuning parameters. The mean (background-corrected) population firing rate is 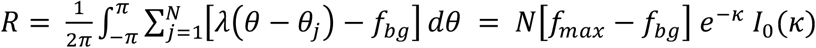) where *I*_0_(*κ*) is the modified Bessel function of first kind. Using 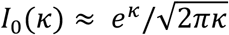, one obtains (Zhang, 1996)

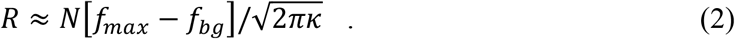

The analytic results obtained for homogeneous populations serve as a good description, also in the case of the related inhomogeneous populations (especially as *N* → ∞, see Results section).

For a description which is as close as possible to the biological findings, we simulate inhomogeneous populations. In such populations, every neuron has individual tuning parameters (*f*_*max,i*_, *f*_*bg,i*_, *κ*_*i*_ and *τ*_*i*_), as shown by the example in Figure 3. Each of these parameters is sampled from a beta distribution, i.e., a probability density function *f*(*x*) ∝ (*x* − *x*_*min*_)^*a*-1^(*x*_*max*_ − *x*)^*b*-1^ on the interval [*x*_*min*_, *x*_*max*_], whose parameters *a* and *b* are obtained from fitting the distribution’s mean and variance to the values shown in Table 1 (see also Figure 2). By shuffling the background firing rates, we ensure that the ratio between a cell’s maximal firing rate *f*_*max*_, and its background firing rate *f*_*bg,j*_ exceeds 5 for every cell, as suggested by Taube (2007).

This method does not aim at exactly replicating the experimental tuning parameter distributions; we rather use the method to explore the *qualitative* effect of inhomogeneous cell properties on the HD code. Nevertheless, since the first two moments of the experimental distributions as well as the observed parameter ranges are reproduced by the beta distributions, they closely relate to realistic HD populations: A comparison of Figure 2 with Figures 5 and 6 in Taube (1995) shows that the model distributions capture key features of the experimental histograms; however, our distributions have less pronounced peaks, especially for *f*_*max,j*_ and *f*_*bg,j*_. To model the preferred head direction of each cell, an equidistant arrangement (*θ*_*j*_ = *j*·2*π/N* − *π*) is distorted by adding to each *θ*_*j*_ a random shift that is chosen from a Gaussian distribution with mean 0 and standard deviation 2*π*/*N*.

**Figure 5.**
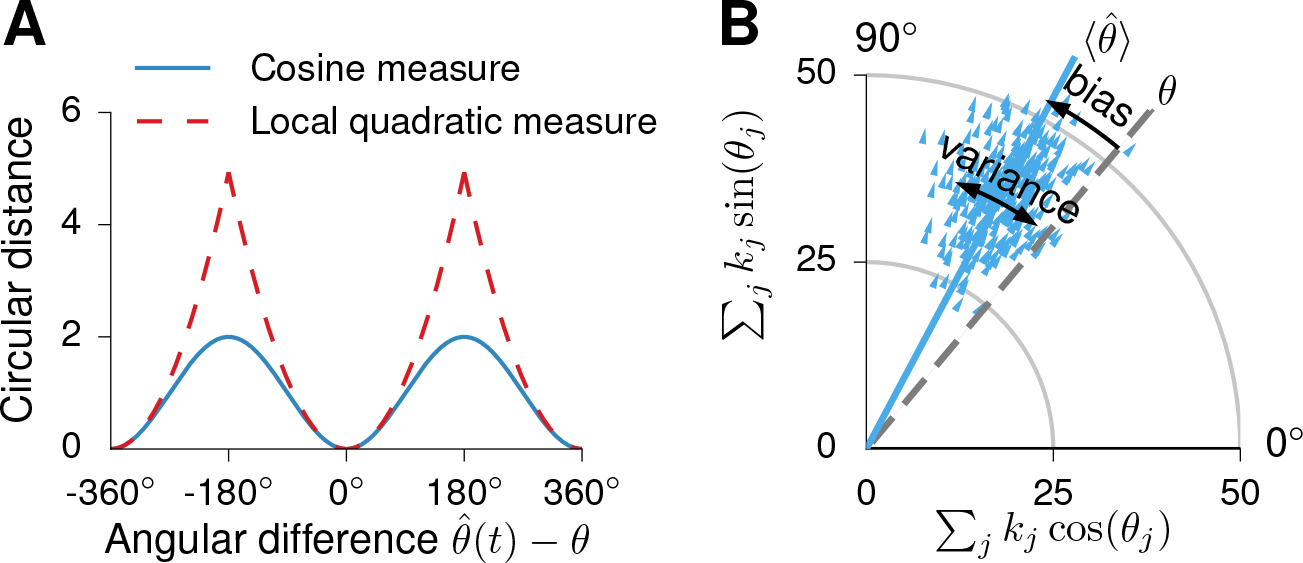
Circular error measure, bias and variance. **(A)** Circular distance function (blue line). For small circular distances this function resembles the squared error (red dashed line). **(B)** 300 realizations of population vectors (blue arrowheads) calculated from a population of *N*=100 neurons encoding a circular static variable *θ*. The mean direction 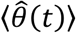 represents the directional average of all possible realizations. If 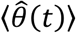 does not agree with the value of *θ* which 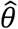 is supposed to represent, as illustrated in the figure, the estimator is called biased. The (circular) variance describes the directional extent of all possible realizations of the population vector given a certain encoded variable (the spread is caused by the noisy encoding).

**Figure 6.**
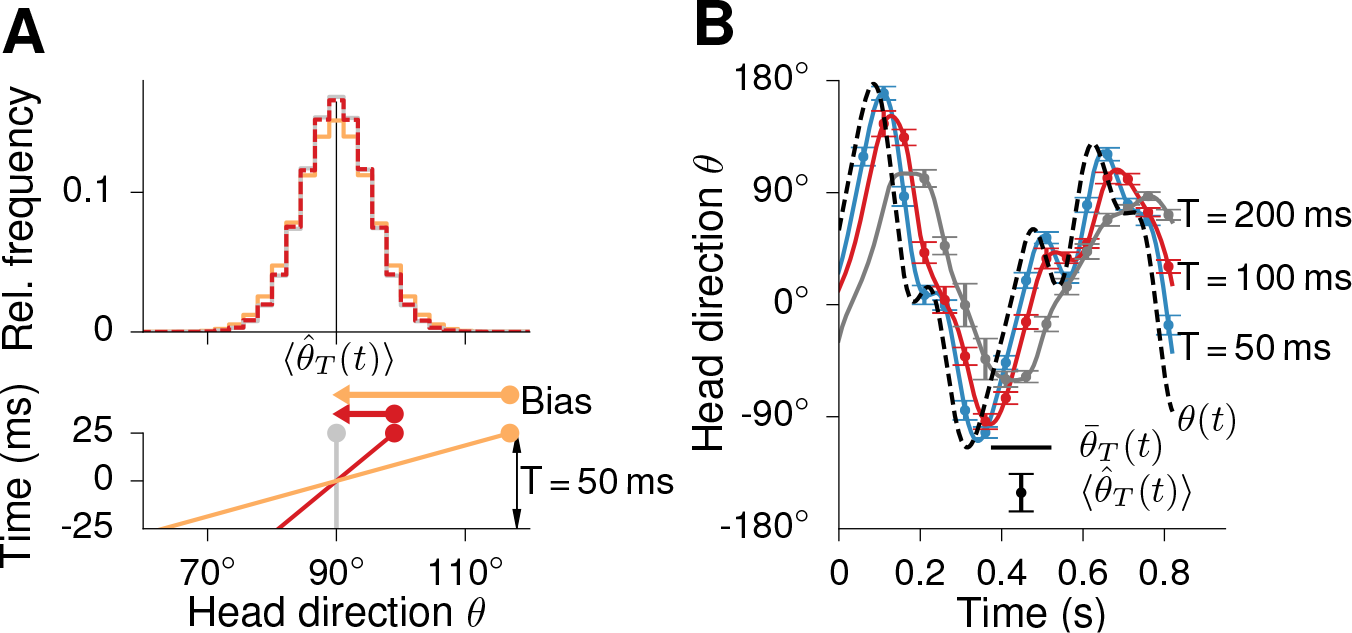
Relation between the population vector and the encoded trajectory. **(A)** Distribution of the population vector’s direction 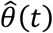 for short trajectory segments with constant angular velocities. The segments are shown below the histogram, their values at *t* = 25 ms are highlighted by large dots. The segment plotted with the red dotted line has *ω* = 360° s^−1^ and represents fast angular head velocities as observed for foraging rats. The threefold faster blue trajectory was chosen to generate a noticeable deviation of the histogram from the static (yellow) case. To sample 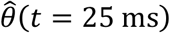, a homogeneous population with *N* = 100 neurons and time-dependent Poisson processes was used to generate spike-count vectors 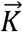 (spike-count window *T* = 50 ms). Even for fast head turns the distribution of 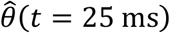 is very similar to the distribution for the static case. In particular, the preferred direction for all three trajectory segments is at 90°, the trajectory average 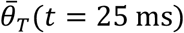, and the variance of the distribution increases only slightly with increasing *ω*. Regarding the trajectory end points, the estimator 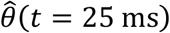 is thus biased for *ω* ≠ 0. **(B)** In essence, the population vector provides a smoothed and delayed copy of the trajectory. The dashed black line denotes an artificial trajectory *θ*(*t*), the continuous lines depict trajectory averages 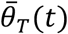 for various window lengths *T*. The mean direction of the population vector agrees well with the time-lagged trajectory averages, i.e., 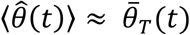 even for trajectory segments with time-varying *ω*(*t*). The mean directions 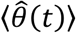 are shown as dots with error bars indicating the standard deviation of the distribution from which they were computed.

### Circular error measure

The accuracy of the HD compass is determined by taking the angle 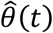 of the population vector (see equation 17) and computing its average deviation from the true trajectory *θ*(*t*). Because we are calculating a difference of circular directions, which are defined on the interval [-π,+ π), measures such as the squared difference between 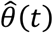 and *θ*(*t*) are not appropriate. Consider, e.g., cases in which 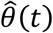 and *θ*(*t*) represent similar directions but lie at different sides of the wrap-around at ±180°, e.g., 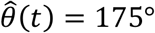 and *θ*(*t*) = − 175°. While the two directions actually differ by only 10°, the squared difference measure would be (350°)^2^. A well-defined error measure for this situation is the circular distance given by 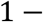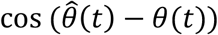, as discussed by Jammalamadaka and Sengupta (2001). For small angular deviations, the circular distance approaches the squared difference, as illustrated in Figure 5A,

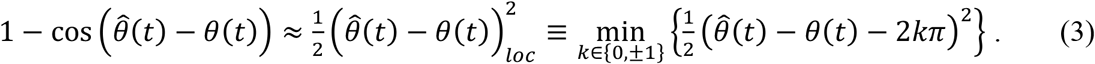

Averaging the circular distance between the population vector’s direction and the true trajectory value over all possible neural responses defines the mean circular error,

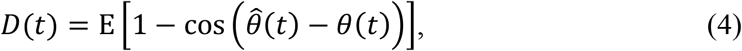

where the expectation is taken over all realizations of the spike-count vector. The mean circular error is the circular equivalent of the mean squared error used for non-directional variables. Averaging the mean circular error *D*(*t*) along the underlying trajectory *θ*(*t*) results in the average mean circular error:

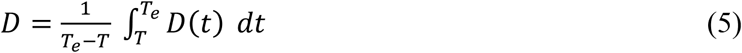

The error measure *D* includes the influences of both the stimulus dynamics and the noisy spike trains on the accuracy of the HD code. Compared to studies on the encoding and decoding of static stimuli, such as Mathis et al. (2012), the trajectory takes the role of the prior stimulus distribution. For graphical representation, we transform the error *D* to the degree-scale by defining the estimator’s accuracy *A* = (180°/*π*) ∙ arccos(1 − *D*), similar to the root mean square error used to describe the performance when dealing with non-circular stimuli.

### Properties of the circular error measure

Because the reader might not be familiar with directional statistics, we briefly introduce the equivalents of the mean value, variance, and bias (see, e.g., Mardia and Jupp, 2009). The equivalent of the mean value is the mean or “expected” direction 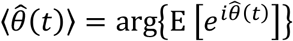 Note that the arg operation is carried out *after* the expectation E is taken, again to deal with the circular wrap-around in a well-defined manner.

The circular variance is given as 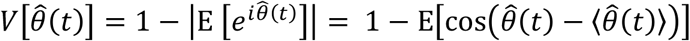 Accordingly, the mean circular error can be decomposed into contributions of the circular variance and a circular bias,

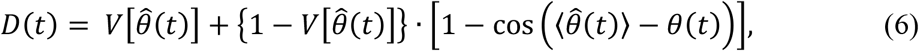

where the second term captures the systematic deviation of the estimator’s mean direction from the actual trajectory value. This latter term is thus the circular equivalent of a bias (see Figure 5 for a visualization of the two error sources) and Equation (6) is the circular equivalent of the decomposition of the mean square error into variance and squared bias.

### Treatment of the zero spike-count case

If not a single HD cell fires in the interval [*t*−*T*, *t*], i.e., 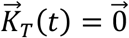, the direction of the population vector is undefined and no information about the current HD is available. The probability for this case can be approximated by 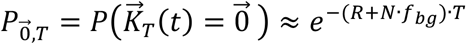 where the population firing rate *R* is defined in Equation (2). To consistently extend the decoding scheme to this case, 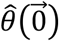 should be picked randomly from *U*[−π,*π*), the uniform distribution on the interval [−*π*, *π*). Separating this special case from all 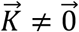, one obtains

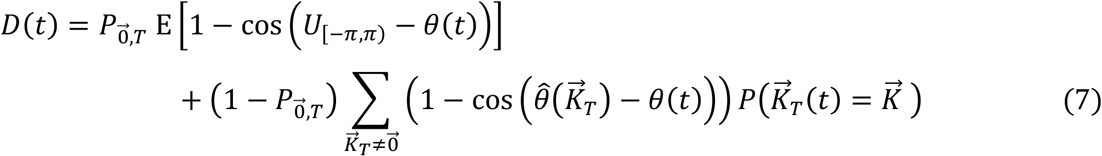

The size of the term 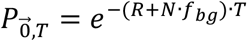 determines whether the case 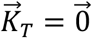 is relevant or not. For the mean parameters of the ADN HD system shown in Table 1, *T*~1/(10 *Hz* ∗ *N*) in order for the zero-spike case to contribute significantly. Even for a small population of only *N* = 100 neurons, this time scale is not more than 1 ms, much smaller than the optimal read-out times, in which we are interested. This shows that the zero-spike case is not relevant for our study so that we will not include the extended definition of the error in our analytical calculations.

### Causal population-vector code

Equation (19) in the main text describes the relation 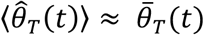 between the expected head direction of the population vector and the corresponding time-averaged true head direction (see Results for the mathematical definition of these quantities). We now derive this relation for two different scenarios; both require homogeneous populations, but for large enough *N* the results are also useful to describe inhomogeneous populations. In the context of 2D place fields and a maximum likelihood decoder, Mosheiff et al. (2017) recently derived a similar relationship.

Let us first assume that the encoded trajectory segments have constant angular head velocity *ω*. This allows us to exploit a symmetry that arises for trajectory segments that are centered around the preferred direction of one particular neuron. For notational simplicity, let us assign the index 0 to this neuron and *θ*_0_ = 0 so that 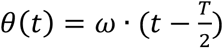 with *t* ∈ [0, *T*]. The other neurons are indexed such that *θ*_−*j*_ = −*θ*_*j*_. This arrangement implies that the activity of the population is symmetric around *θ* = 0 in the time interval *t* ∈ [0, *T*], i.e. *α*_−*j*_(*t* = *T*) = *α*_*j*_(*t* = *T*). Introducing the notation 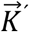 for a count vector with mirror indices relative to 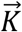 i.e., 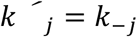, the symmetric activation implies 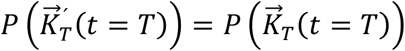, while 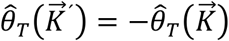. Using these identities the mean direction 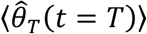 of the estimator 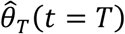 can be calculated as

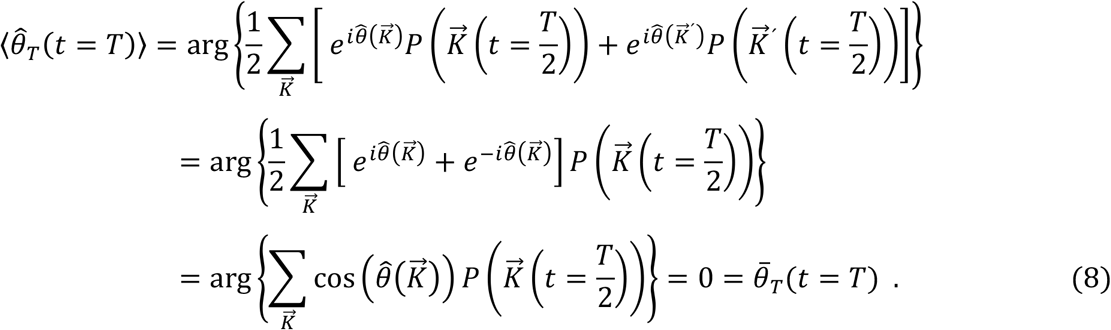

An illustration of the symmetry of the distribution of 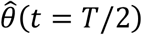 around the trajectory average of a trajectory segment with constant angular velocity is shown in Figure 6A.

For a static encoded variable, the mean population vector can be calculated as done by Glasius et al. (1997). Applying their approach to a general, time-varying head-direction signal *θ*(*t*), we obtain

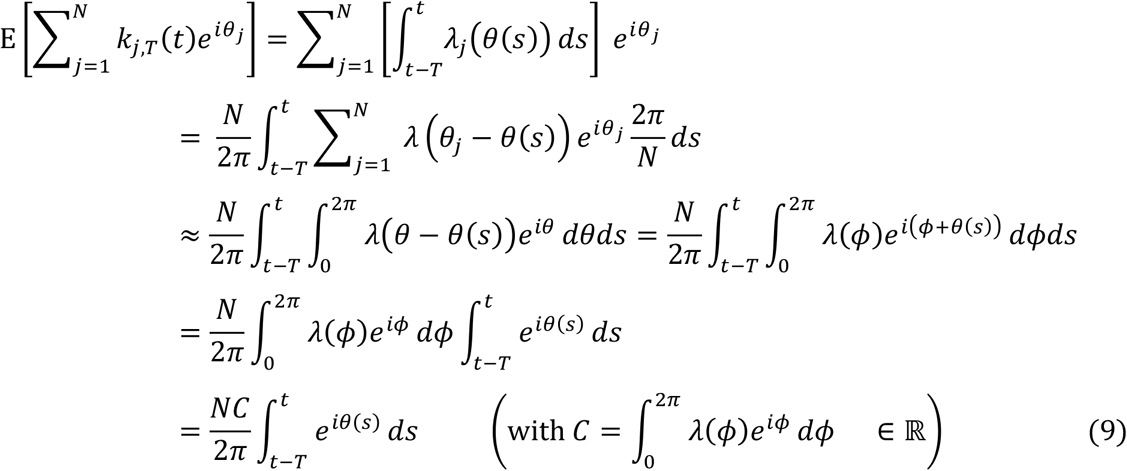

so that

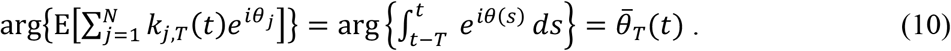

Thus, the mean population vector points into the direction of the trajectory average 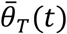. We cannot directly transfer this result to the mean direction 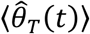 of the population vector, but it is reasonable to expect a similar behavior. Indeed, as shown by the example trajectory in Figure 6B, the numerical estimates agree well with 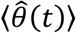. As also demonstrated in this figure the population vector effectively smoothens a trajectory and delays it by a time lag of *T*/2.

The histogram in Figure 6A shows that for reasonable head-direction velocities and neural population sizes, the distribution of 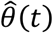 is narrow and closely concentrated around 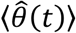. We may thus expand the mean circular error *D*(*t*) for small local angular distances to first non-vanishing order as 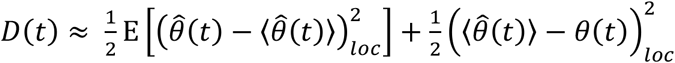. To obtain an approximation of the time-averaged circular error *D*, we integrate this expression along the whole trajectory over the corresponding time window, i.e., *T*_*e*_ − *T* where *T*_*e*_ denotes the trajectory’s duration,

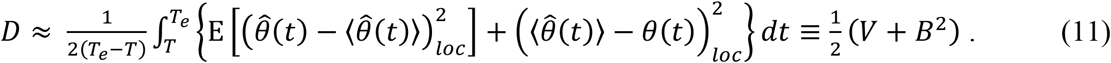

The symbols *V* and *B*^2^ denote the (approximate) time-averaged variance and time-averaged squared bias, respectively.

Figure 6 also shows that the width of the 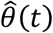 distribution is only mildly affected by the angular head velocity, even for high velocities. This implies that its variance *V* depends only weakly on time *t* so that we can omit the temporal average when calculating *V*. Furthermore, as the 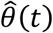 distribution does hardly vary with *ω*, results by Seung and Sompolinsky (1993) for the (local) variance of the direction of the population vector (derived assuming a static stimulus) apply. Their derivation is based on Fisher information. We can use it to approximate the local squared deviation of the population vectors angle from its mean direction as

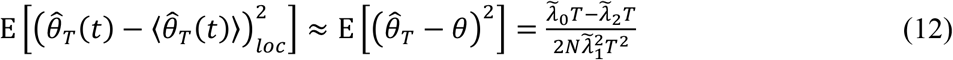

where *θ* is a static stimulus parameter, 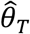 its estimator, and 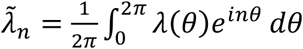 are the Fourier components of the zero-centered tuning curve *λ*(*θ*). For von Mises curves (2), one obtains 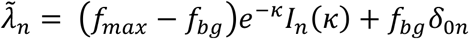 since 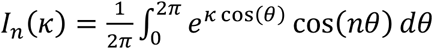. Inserting the first two terms of the expansion of each Bessel function for large *κ*, we arrive at

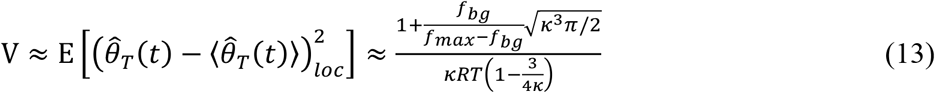

for a homogeneous neural ensemble with von Mises tuning curves (see Figure 7, dotted lines). This shows that as the average number *RT* of (non-background) spikes in the spike-count window increases, the variability *V* decreases and more information about the stimulus is available. Similarly, the variance of the estimator increases for higher background activity (Seung and Sompolinsky, 1993; Mathis et al., 2012).

**Figure 7.**
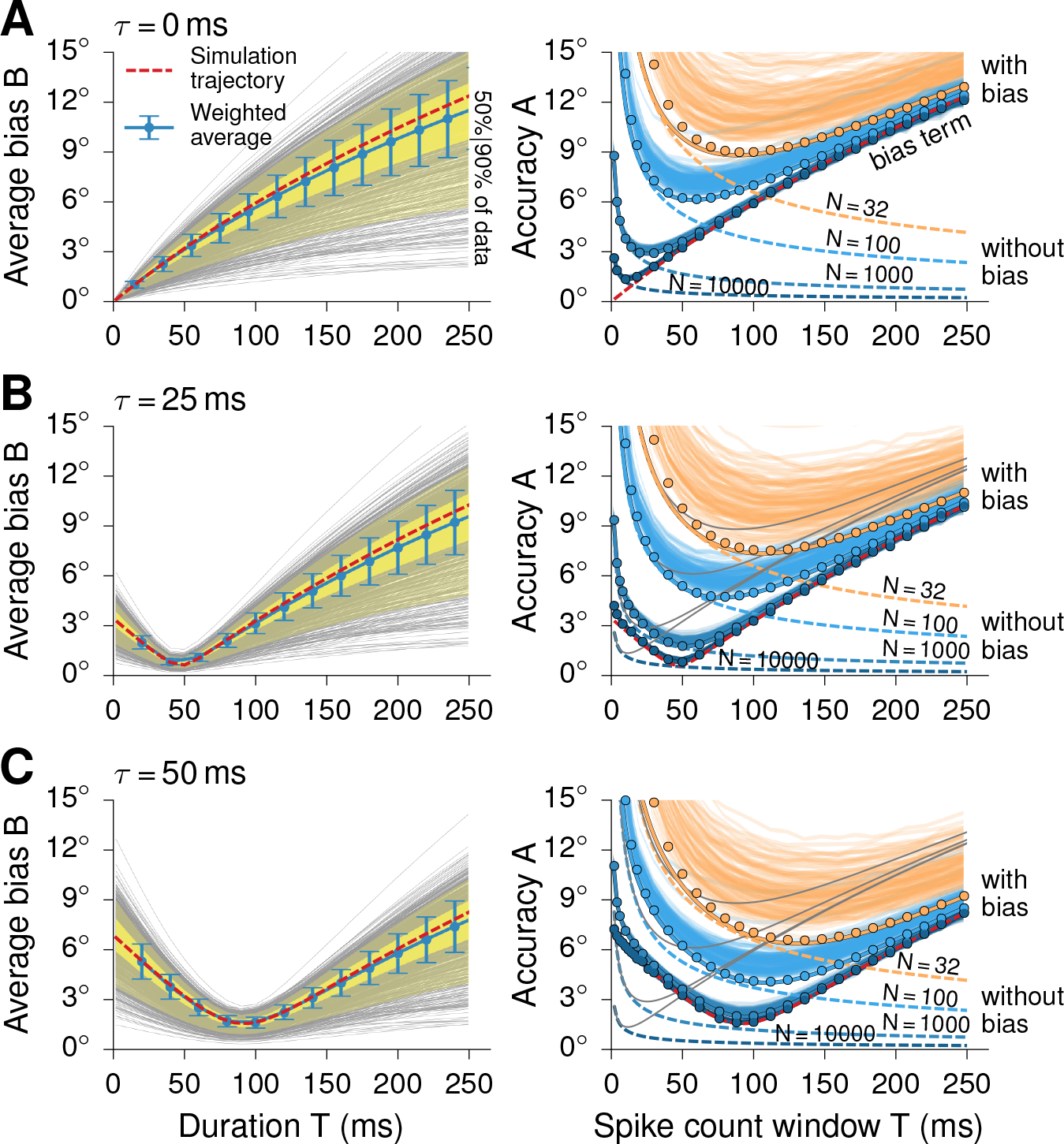
Bias and accuracy of the population vector. **(A, left panel**) Mean bias *B* for all trajectory segments of the data set (Mizuseki et al., 2009) as a function of segment duration *T*. The duration-weighted average of all trajectory segments (with error bars indicating the duration-weighted standard deviation) is shown as well. Light and darker yellow areas indicate the sets of trajectory segments between the 25th–75th and 5th–95th percentiles, respectively. The red dashed line corresponds to the representative trajectory which was used as a basis of the simulations. For causal decoding without anticipation, the plotted data represent the best achievable accuracy for given *T*. Since longer time windows admit stronger changes of the head direction, the bias term increases with increasing *T*. **(A, right panel)** Approximation results for the reconstruction accuracy given various population sizes (continuous lines). For each population size there is a spike-count window which leads to an optimal accuracy. Larger populations can reach a higher HD accuracy. While the contribution of the variance to the total error decreases with the size of the population (dashed lines), the bias term (red dashed line) is independent of the population size. Dots represent simulation results from Monte Carlo simulations using homogeneous populations. The transparent lines are simulations results from different inhomogeneous populations (one line per population). As evident from the figure, the approximate solution derived in the text captures the population vector accuracy of homogeneous populations as well as of inhomogeneous populations (for large enough population size). **(B,C)** As in (A) but for anticipatory time intervals τ = 25 ms and τ = 50 ms, respectively. Left panels: In both cases, the dip with smallest average bias B is found at T ≈ 2τ because for these values the interpolated trajectory φ(t) is closest to θ(t), on average. The length of the spike-count window with minimum bias is thus controlled by τ. Right panels: In addition to data for the nonzero τ, the analytic results from (A) are plotted as gray background lines. The simulation results (dots and transparent lines) again agree well with the analytic description (continuous, nontransparent lines), especially for large population sizes N. As the bias term has a minimum at T > 0, longer spike-count windows can be used compared to the non-anticipatory case. This reduces the variance and bias of the estimator and results in smaller minimal reconstruction errors. However, too long ATIs can also deteriorate population codes. One such example is shown for which the optimal accuracy of a population with N = 10000 neurons is worse for anticipatory firing with τ = 50 ms than for τ = 25 ms.

To first approximation, the time-averaged circular error can be written as

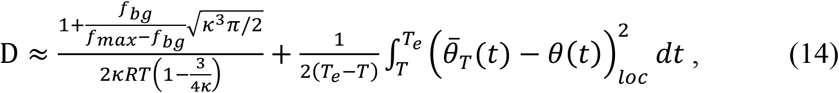

where the second term needs to be evaluated for the encoded trajectory *θ*(*t*). In the right column of Figure 7, Equation (14) is shown as a function of *T* using the representative trajectory segment *θ*(*t*) mentioned above. The tuning curve parameters are chosen according to the mean values shown in Table 1.

### Monte Carlo simulations

We use Monte Carlo simulations of the average circular error to check the analytical results and assess the effect of inhomogeneous tuning parameters. To this end, we sample the integral over the trajectory and the Poisson noise for *M* ≫ 1 times: first, an endpoint of a trajectory segment *t*_*i*_ ∈ [*T*, *T*_*e*_] is randomly selected. Then, a count vector 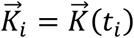 is sampled from the respective Poisson distribution. Finally, the circular distance 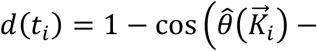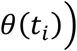 is calculated and the average circular error is approximated as 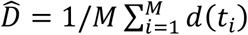.

### Experimental design and statistical analysis

We reanalyzed data originally recorded by Mizuseki et al. (2009) and refer the reader to that publication for details on the experimental design. All our analyses were performed in Python (RRID:SCR_008394) using functions from the Scipy (RRID:SCR_008058) and Numpy (RRID:SCR_008633) modules.

## Results

The head-direction (HD) system of rodents encodes the time-dependent azimuth *θ*(*t*) of the animal’s head relative to a landmark-anchored reference direction (Figure 1) and plays a functional role for spatial navigation (Butler et al., 2017). Notably, during head turns, HD cells start to fire before their preferred direction *θ*_*j*_ is reached, as though the neurons anticipated the future head direction (Blair and Sharp, 1995). Faster rotations lead to stronger shifts in time than slow turns. Denoting the time-dependent angular head velocity by *ω*(*t*), the shift can be approximated by

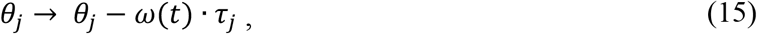

where the time constant *τ*_*j*_ is called the anticipatory time interval (ATI) of cell *j* and describes the extent to which HD firing seems to be shifted forwards in time (Blair and Sharp, 1995; Blair et al., 1997; Taube and Muller, 1998; see the right panel in Figure 2A for a visualization).

The rodent HD system consists of multiple processing stages along which ATIs decrease from stage to stage: from the lateral mammillary nuclei (*τ* ≈ 40-75 ms) to the anterodorsal thalamic nucleus (ADN) and retrosplenial cortex (*τ* ≈ 25 ms) to the postsubiculum, where neurons tend to neither lag nor anticipate the cell’s preferred firing direction (Taube 2007). As with other HD cell parameters (see also Table 1), ATIs vary from cell to cell (Blair et al., 1997; Taube and Muller, 1998). At the mechanistic level, ATIs have been explained by an interplay between afferent dynamics and movement statistics (van der Meer et al., 2007; Tsanov et al., 2014; see also Zhang, 1996) but a consistent functional interpretation of ATIs is still missing.

As we will show, anticipatory firing improves the accuracy with which the *present* head direction can be decoded from time-dependent HD signals. For concreteness, we use the population vector decoder (Georgopoulos et al., 1983, 1986), which provides a reliable estimate of variables encoded by the distributed activity pattern of a neural ensemble (Seung and Sompolinsky, 1993; Salinas and Abbott, 1994; Glasius et al., 1997; Stemmler et al., 2015). We focus on HD cells in the ADN, which is a core region of the HD system (Taube, 2007) and contains the highest fraction of HD cells (Taube, 2003). To cover the dynamic character of head movements we explicitly consider time-varying stimuli (Mosheiff et al., 2017). We take a computational approach and use experimentally recorded HD trajectories *θ*(*t*) to generate artificial spike trains (Figure 3). The periodic tuning curves of individual HD cells can be captured by von Mises functions (Zhang et al., 1996) whose tuning width, peak firing rate and background firing rate are chosen according to the available experimental literature (Figure 2B). Spike generation is modeled by inhomogeneous Poisson processes, see Equation (1). From the spike trains of this simulated neural population, we compute an estimated trajectory 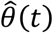 based on the population-vector decoder. We then measure the accuracy of the HD system by analyzing the distribution of 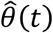 and comparing its mean 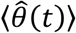 with the true trajectory *θ*(*t*), as described in Materials and Methods. Using this framework, we systematically explore the HD representation and characterize the effect of anticipatory firing. We derive an analytic approximation for the population code’s accuracy and use numerical simulations to check its validity and to study populations with inhomogeneous tuning curves. An approach based on a reconstruction from experimentally measured spike trains, as in Johnson et al. (2005) for the postsubiculum, is currently not feasible because simultaneous recordings of HD cells in ADN are only available for small populations of less than 20 HD neurons (Peyrache et al., 2015; Taube, 2015).

### Head-direction trajectories

The head-direction trajectories *θ*(*t*) for this study are taken from a data set which was recorded in the lab of G. Buzsáki (Mizuseki et al., 2009). Since the head direction is a circular variable, temporal averages and expectation values need to be consistent with circular statistics (see Materials and Methods). In particular, the temporal average 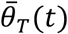 of a trajectory segment θ(*s*) within some time interval [*t* − *T*, *t*], is defined by

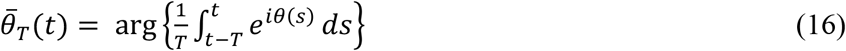

so that for constant angular head velocity, i.e., *θ*(*t*) = *θ*_0_ + *ω* ∙ *t*, the trajectory average corresponds to the midpoint of the trajectory segment, 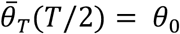. To avoid overly complicated mathematical expressions, we will omit indices, such as *T* in 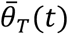, or variables, such as *t* in 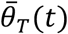, unless this could cause misunderstanding.

### Tuning curves and spike-train model

Head-direction cells in the ADN have been studied in great detail (see, e.g., Blair and Sharp, 1995; Goodrich and Taube, 1997; Blair et al., 1997, 1998; Stackman and Taube, 1998; Taube and Muller, 1998; Taube, 2007; Clark et al., 2009, 2012). Their stereotypical tuning profiles show a single hump and are well fit by von Mises functions (Zhang et al., 1998),

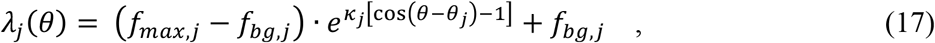

with peak firing rate *f*_*max,j*_, background firing rate *f*_*bg,j*_, “concentration” parameter *κ*_*j*_, and preferred direction *θ*_*j*_. The cell’s tuning width *σ*_*j*_ can be calculated as 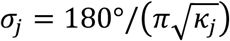.

Based on these tuning curves each model neuron generates an inhomogeneous Poisson spike train, see Equation (1), with time-dependent rate *λ*_*j*_ (*θ*(*t*)). If we use *λ* without subscript, we refer to a neuron with preferred direction *θ* = 0.

According to Taube and Bassett (2003), there are around *N* ≈ 10,000 HD cells in ADN per brain hemisphere. We therefore study ensembles up to this size but also consider smaller *N* to analyze size-dependent effects.

The tuning parameters of HD cells vary strongly from neuron to neuron. To evaluate the effect of such variations on the neural code we describe the parameters by probability distributions whose ranges, means, and standard deviations are estimated from the literature (see Table 1, as well as Materials and Methods). Sampling the parameters of each model neuron from these distributions results in what we call an inhomogeneous population (see Figure 3). To model the preferred head direction of each cell, the equidistant arrangement (*θ*_*j*_ = *j*·2*π/N* − *π*) is altered by adding to each *θ*_*j*_ a random shift chosen from a Gaussian distribution with mean 0 and standard deviation 2*π*/*N*. The mathematical analysis focuses on homogeneous populations. Here, neurons differ only by their (equidistant) preferred direction but share all other parameters (so that we omit the subscript *j*), chosen as the mean values in Table 1. For simulations of inhomogeneous populations, we sample ATIs from the distribution shown in Figure 2; for homogeneous populations, we assume *τ* = 25 ms for all cells, unless noted otherwise.

### Causal population vector reconstruction

The Population Vector *PV*(*t*) is a weighted sum, 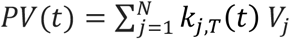, where the *V*_*j*_ are unit vectors in *θ*_*j*_-direction and *k*_*j,T*_(*t*) denotes the number of spikes fired between time *t* − *T* and *t* (Georgopolous et al. 1983, 1986). Introducing the phasors exp(*iθ*_*j*_) = cos(*θ*_*j*_) + *i* sin(*θ*_*j*_), the population vector can be written as 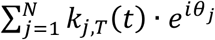 its direction is given by the argument of this complex number. Defining the spike-count vector 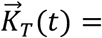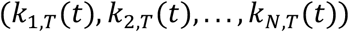, the animal’s head direction can be decoded from the experimental or modeled HD cell responses as

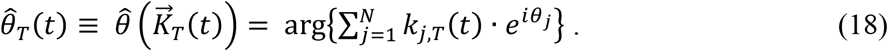

Note that the decoder 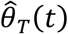 is chosen to obey causality: only spikes from spike-count windows ending at time *t* are used to estimate *θ*(*t*), as illustrated in Figure 4. This choice reflects the fact that any downstream processing step of the head direction signal can only depend on spikes that were fired in the past.

### Read-out accuracy: a function of systematic lag (bias) and random noise (variance)

The theoretical framework now at hand allows us to calculate how accurately the current head direction *θ*(*t*) can be determined from the time-dependent population activity of non-anticipatory HD cells in a moving animal. Since the PV estimator 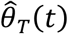 relies exclusively on spikes fired before time t, the computed head direction always lags behind the current head direction *θ*(*t*). In statistical terms this means that 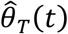 is *biased* with respect to *θ*(*t*). Additionally, since the spiking behavior of an ensemble of neurons is noisy, 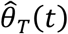 exhibits random fluctuations, which are measured by the variance *V*. As shown in Materials and Methods, the (circular) error *D* for the HD read-out from a neural ensemble is given by the sum of the (squared) bias *B* and the variance *V*,

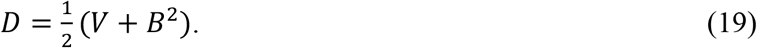

In the following, we will describe the behavior of the bias *B* and the variance *V* of the causal PV estimator and demonstrate at which point anticipatory firing introduces an improvement of the read-out accuracy.

To determine the bias *B* of the PV estimator, we start by evaluating the estimator’s average at a given instance of time. As derived in Materials and Methods,

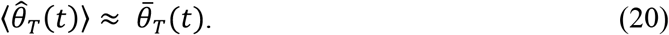

This means that when averaging out the noise induced by stochastic spiking, the PV estimator provides the average HD of an animal during the spike-count time *T*. Since the bias is given by the difference between 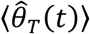 and *θ*(*t*), the dynamic nature of the HD induces a systematic error. To visualize this, consider a trajectory segment with constant AHV: *θ*(*t*) = *θ*_0_ + *ωt*. Here 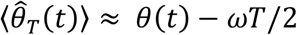, such that the bias amounts to *B* = *ωT*/2. The PV estimator is thus biased for every spike-count window for which the average HD does not match the final HD. Generally, longer spike count windows will lead to larger biases, because the HD has more time to change so that the average HD deviates stronger from the endpoint-HD. We evaluated the average squared bias *B*^2^ along all experimentally recorded trajectories as a function of *T*. The results are shown in Fig. 7A and all share an increase with integration time *T*.

Based on this analysis of the bias, it appears reasonable to use very short spike count times to keep the spikes used for decoding as close as possible to the current time point *t*. This will, however, increase the noise of the read-out as less and less spikes are involved in the estimation and their stochastic variations cannot be averaged out anymore. Together, the two opposing effects define an optimal read-out time for which there is an ideal trade-off between systematic lag and statistical fluctuations.

The amount of error induced by noisy spiking (i.e., the variance *V*) can be approximated analytically using Fisher information. Since this spiking noise is independent of the AHV, we can refer to results derived for the well-studied static case (see Materials and Methods, Equation 13, for the full result). We find that the variance decreases with the average number of collected spikes as 1/RT, thus decreasing as spikes are collected during longer time windows (see Figure 7A, dotted lines).

The trade-off arising from the two different error sources is visualized in the right column of Figure 7. Along with data points obtained from applying our analytical treatment to an experimentally recorded trajectory, we show results from Monte Carlo simulations (see Materials and Methods). These involved either homogeneous or inhomogeneous HD cell ensembles and confirm our mathematical prediction. In particular, as *N* increases, results from inhomogeneous and homogeneous cell ensembles align since all possible HDs are uniformly covered by tuning curves.

When decoding time-independent stimuli, the read-out window *T* is usually assumed to be a free parameter that needs to be chosen in a biologically or methodologically useful way (see e.g. Seung and Sompolinsky, 1993; Brown et al., 1998; Zhang and Sejnowski, 1999; Mathis et al., 2012). Considering the dynamic nature of a stimulus, as we do in this study, leads to specific spike-count windows *T* for which the highest possible accuracy is achieved (see also Mosheiff et al., 2017). Note that the bias depends exclusively on the shape of the HD trajectory, while the variance depends exclusively on the parameters of the neural ensemble. For a *non-anticipator*y population of HD cells, the only chance to decrease the optimal read-out error is to invest more neural resources (*N*, *f*_*max*_) and thus reduce the contribution of the random spiking noise to the error. The systematic lag can, however, not be changed directly. We will show in the following paragraph that anticipatory firing does influence the bias term and can lead to significant improvements of the HD code.

### Effect of anticipatory firing in HD ensembles

As proposed by Blair and Sharp (1995), we assume that the shift of the preferred head direction scales linearly with angular head velocity. Writing 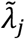 for the average firing rate of an anticipatory neuron in a homogeneous population and inserting Equation (15) for the preferred direction, we obtain

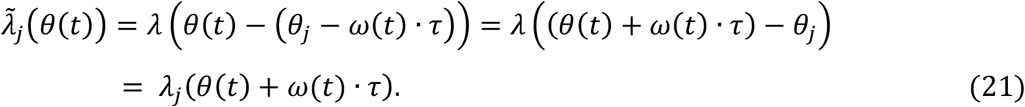

This formulation treats anticipation in HD cells as a shift in the angular trajectory given by *φ*(*t*) = *θ*(*t*) + *ω*(*t*) ∙ *τ*, as illustrated in Figure 8. If the AHV *ω*(*t*) were constant between *t* and *t* + *τ*, the product *ω*(*t*)*τ* equals the total turning angle in that time interval.

**Figure 8.**
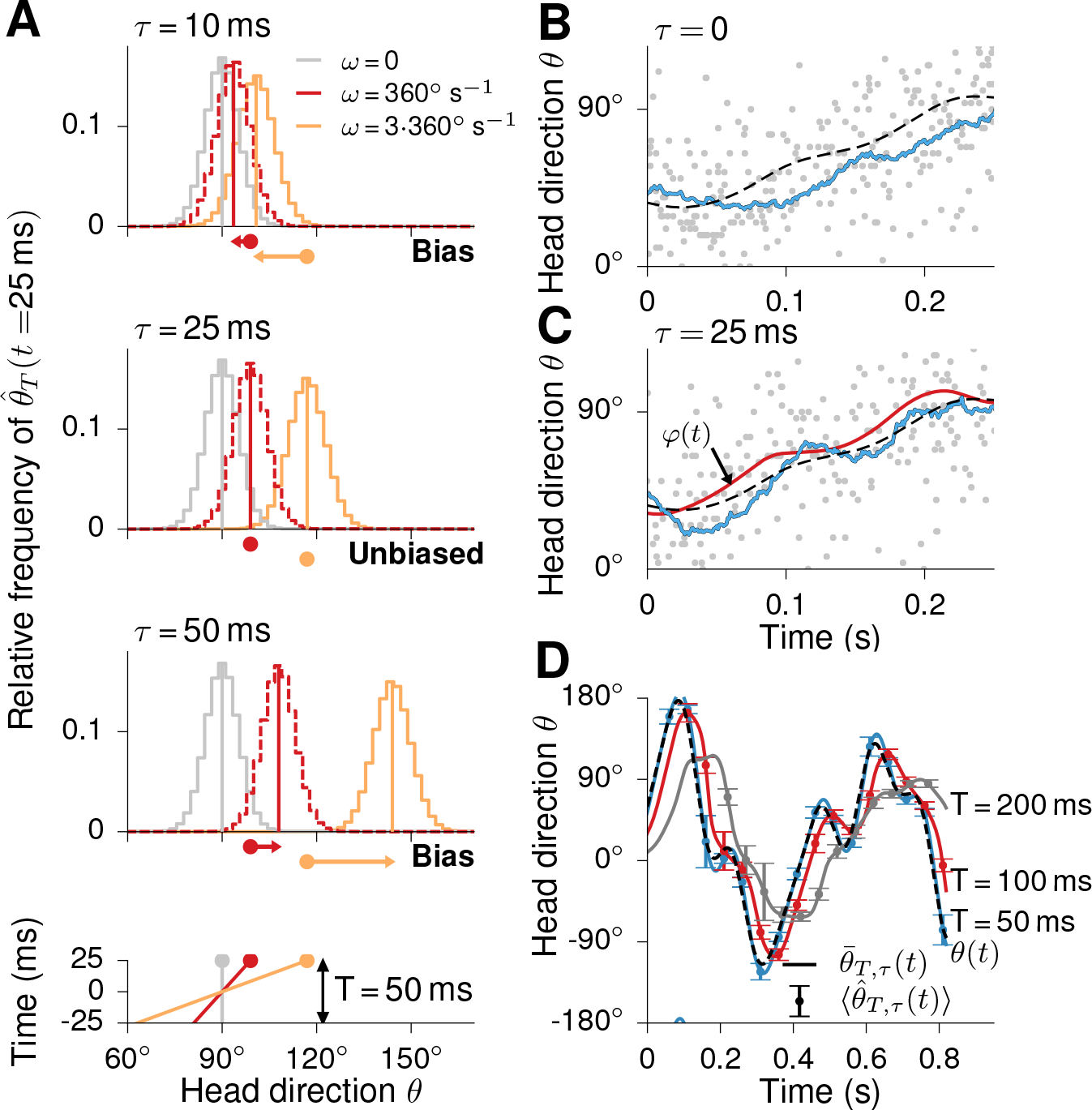
Effects of anticipatory firing on population-vector decoding. Effect of anticipatory firing on the distribution of 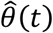 and its bias. **(A, bottom row)** The three trajectory segments with constant AHVs ω (0, 360°/s, 3·360°/s) used for the simulations (as in Figure 6). **(A, three upper rows)** Distribution of 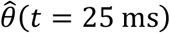 for τ = 10*ms*, τ = 25*ms* and τ = 50*ms*, respectively. An anticipatory time interval (ATI) of τ = 10*ms* leads to a reduction of the bias compared to the non-anticipatory case shown in Figure 6. The bias is reduced as the expected directions 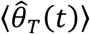, denoted by the vertical lines, are closer to the trajectory’s end points (depicted as dots). For τ = 25*ms* the condition τ = *T*/2 (see main text) is fulfilled and the expected directions agree exactly with the trajectory’s end points. In this situation 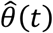 is an unbiased estimator of *θ*(*t*). For τ = 50*ms* the bias is as large as in the non-anticipatory case, but with opposite orientation. Note that the width of the distributions does not depend on the ATI. **(B)** Raster plots of 100 HD cells encoding the trajectory segment *θ*(*t*), depicted as black dashed line, without anticipation (τ = 0). The decoded trajectory 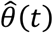 is shown as blue line and lags behind the true trajectory *θ*(*t*). **(C)** With an anticipation of τ = 25*ms*, the decoded trajectory does no longer lag behind the true trajectory. The red line illustrates the modified trajectory *φ*(*t*) = *θ*(*t*) + *ω*(*t*)*τ*. **(D)** Same as in Figure 6 but for anticipatory firing with τ = 25*ms*. The lagging circular averages of the modified trajectory *φ*(*t*) are shown as continuous lines. They agree well with the mean directions 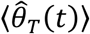 which were computed from the anticipatory spike trains.

Accordingly, the expected direction of a homogeneous population of anticipatory neurons is given by

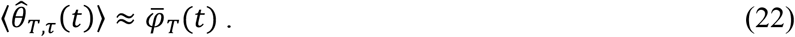

The subscript τ indicates that we are describing an anticipatory HD population. Since the trajectory *φ*(*t*) corresponds to an estimate of the future HD, this change of the expected direction 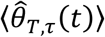 can lead to a compensation of the bias of 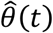 compared to the non-anticipatory case (equation 20).

To better understand this effect, we first examine trajectory segments with constant AHV *ω*, i.e., with *θ*(*t*) = *θ*_0_ + *ω* ∙ *t*. In this case *φ*(*t*) = *θ*_0_ + *ω* ∙ (*t* + *τ*) = *θ*(*t* + *τ*) so that *φ*(*t*) is a time-shifted version of *θ*(*t*). We obtain 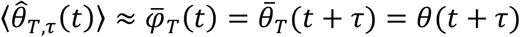 for the expected direction of the causal estimator 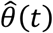. The ATI *τ* thus temporally shifts the expected direction of 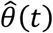 along the trajectory *θ*(*t*). Evaluation of the average bias term for a linear trajectory segment leads to *B* = |*ω* ∙(*T*/2 − *τ*)|. Choosing *τ* within (0, *T*) reduces the bias compared to the situation without anticipation. For constant AHV, it is even possible to fully remove the bias by choosing *τ* = *T*/2, as illustrated in Figure 8.

Anticipation also reduces the bias when the velocity of the HD *θ*(*t*) varies in time. To quantify this effect in the experimentally relevant case, we re-calculate *B* as for Figure 7A but with anticipatory firing. We use Equation (21) to compute the expected HD. The results of this analysis are shown in the left panels of Figure 7B and 7C for *τ* = 25 *ms* and *τ* = 50 *ms*, respectively. In stark contrast to populations without anticipation (Figure 7A), *B* does not increase monotonically with *T* but has a minimum. Similar to the case of trajectories with constant AHV, the average bias *B* is smallest for *T* ≈ 2*τ*. In this case, the implicit interpolation into the future described by *φ*(*t*) has the optimal size. However, because of the time-varying nature of the AHV, the bias is not completely canceled.

In contrast, the variance of 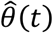 does not depend on the anticipatory time interval *τ*, as seen by considering trajectory segments with constant AHV or the data shown in Figure 8. To describe the reconstruction error *D* for anticipatory firing, we directly adopt the variance term from Equation (13) and replace the bias term by the anticipatory bias as analyzed above. We now numerically calculate the average circular error *D* with homogeneous and inhomogeneous populations. For the inhomogeneous populations, the ATIs are sampled from the distribution depicted in Figure 2 (for *τ* = 50 *ms*, the distribution of *τ* is shifted by 25 *ms* to the right). The right panels of Figure 7B and 7C show the analytical result from Equation (13) together with simulation results for ATIs of *τ* = 25 *ms* (to mimic experimental ATIs) and *τ* = 50 *ms* (to illustrate the effect of longer anticipation).

As in Figure 7A, the data shown in Figure 7B and 7C demonstrate a qualitative agreement between the analytic description and the simulations. Most notably, the results for populations with realistic ATI distribution align closely with the results for homogeneous populations (for sufficiently large *N*) and show that cell-to-cell ATI variability does not reduce the advantages of anticipatory firing. Compared to populations without anticipation, the minimal errors of anticipatory populations are shifted towards longer spike-count windows *T* because the bias does not increase monotonically with *T* but rather has a minimum for *T* > 0. Consequently, spikes can be collected over longer time windows (as long as the bias remains small) which reduces the variance. For *τ* = 25 *ms*, this improved trade-off leads to a reduction of the minimal error for every population size shown in Figure 7B. For large ATIs, however, the minimum achievable bias can become so high that the minimal reconstruction errors are larger than without anticipation (Figure 7C). Regarding the neural code of the causal population vector decoder, anticipation thus only improves accuracy if the ATI lies within a certain interval.

In Figure 9A, we show how much the minimum achievable error can be reduced by anticipatory firing with different ATIs. To generate this figure, the function *A*(*T*) was numerically minimized for *τ* = 0 *ms* and various values *τ* > 0 *ms*. The minimum errors for *τ* = 0 *ms* are shown in the inset as a function of population size *N*. The main plot presents the (average) fraction of the minimum *A* for and without anticipatory firing. The figure reveals that anticipatory firing can lead to large improvements in reading out the HD compass: the experimentally determined average value of *τ* = 25 *ms* increases the accuracy by up to 40%. Figure 9B demonstrates how large the population size *N*_0_ of a non-anticipatory population has to be to reach the same accuracy as an anticipatory population of size *N*. For a large range of population sizes and ATIs, non-anticipatory populations need to be more than three times as large as populations of anticipatory neurons (Figure 9B). For *τ* = 25 *ms*, the ratio *N*_0_/*N* is particularly high for (biologically plausible) population sizes between 1,000 and 10,000 neurons. These results reveal that the experimentally reported value of *τ* = 25 *ms* lies exactly in the range that leads to an optimal improvement of the HD code given the broadly tuned neurons and the temporal statistics of the head movements.

**Figure 9.**
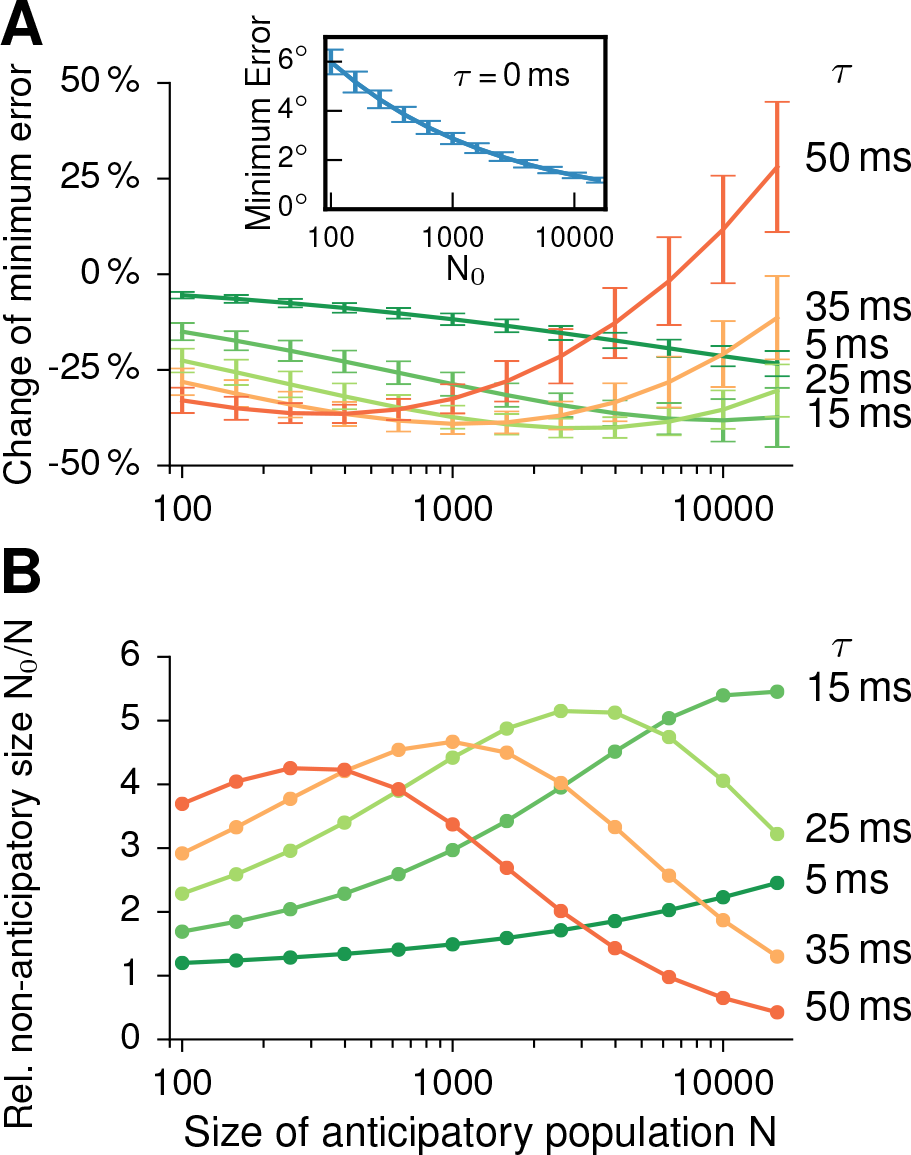
Anticipation improves the accuracy of the head-direction compass. **(A)** The inset shows the best accuracy A(T) achievable for a homogeneous population with non-anticipatory neurons. The main figure depicts the relative change of this smallest error if HD cells fire in an anticipatory manner. Data points are duration-weighted averages over all trajectory segments, error bars denote the duration-weighted standard deviations. For population sizes between 1,000–10,000 neurons, the biologically realistic value τ = 25*ms* belongs to the best options and improves the accuracy by up to around 40 %. Anticipation can, however, also deteriorate the code quality, in our examples for τ = 50*ms* and population sizes N > 6000. **(B)** Increasing the population size reduces the minimum error, too. The figure shows how much larger the population size *N*_0_ of a non-anticipatory population needs to be to yield the same effect as an anticipatory population of size N (color code as in A). For example, the performance of a population with N = 10,000 neurons and an ATI of τ = 25*ms* matches that of a population with N = 40,000 without anticipation.

## Discussion

Single HD cells in the rat anterodorsal thalamic nucleus (ADN) predict the animal’s head direction roughly 25 milliseconds into the future (Blair and Sharp, 1995; Taube and Muller, 1998). Ironically, such anticipatory firing helps to encode the *present* HD, not the future HD, as we have shown here by analyzing the read-out process. Taking the temporal statistics of head movements in foraging rats into account, we found that the error in modeled neuronal population vectors is smallest when HD cells anticipate the true HD by an anticipatory time interval (ATI) of around 25 milliseconds for populations with a few thousand neurons, in agreement with the ADN measurements. With realistic parameters for the HD tuning curves, anticipatory firing decreases the minimal error in the HD estimates by up to 40%. One would need a five-fold increase in the number of neurons to achieve the same performance in the absence of anticipation.

Because of their simplicity, population vectors are often the method of choice to decode the HD signal from calcium imaging data (Peyrache et al., 2015) and multi-electrode recordings (Kim et al., 2017). Moreover, population vectors are readily interpreted in the framework of ring-attractor networks, which posits that vestibular and other sensory signals drive bumps of neuronal activity to move along a ring-like network of neurons, as reviewed in Laurens and Angelaki (2018). Other decoding approaches, such as spike-by-spike population decoding (Huys et al., 2007), Bayesian filtering (Bobrowski et al., 2009) or predictive Kalman filters, are not likely to provide significant advantage over population-vector decoding. Indeed, in a similar decoding task, namely the reconstruction of spatial trajectories from place-cell activity, such alternative strategies did not improve the decoding accuracy (Brown et al., 1998).

A number of questions lie outside the scope of this study. We did not model path integration based on accumulating and error-correcting the HD signal, as proposed by Cheung et al. (2008). Moreover, we did not address the mechanistic origin of anticipatory firing. Possible explanations for anticipatory firing range from neurons sensing head acceleration (Zhang, 1996) to high-pass filtering based on short-term synaptic depression (Puccini et al., 2007) or a combination of spike rate adaptation and post-inhibitory rebound (van der Meer et al., 2007). Instead, we focused on the role anticipatory firing plays in reducing the bias in estimating the head direction signal as it changes in time.

Anticipation resolves a quandary inherent in decoding any time-varying quantity from a population of spiking neurons. Given the discrete and stochastic nature of neuronal discharges, averaging over time should improve the accuracy of decoding, were it not for the fact that the stimulus is continually changing. If the decoder reads out the population activity over a specific time window *T*, a trade-off arises (see, e.g., Brown et al., 1998; Zhang et al., 1998; Mosheiff et al., 2016; and our approach): the dynamics of the stimulus causes the deviations in the population-vector read-out (relative to the true, instantaneous stimulus) to increase with *T*, whereas the variance for estimating a static stimulus would decrease with *T* (Fiete and Burak, 2012). Systematic biases caused by averaging over time are exacerbated by the requirement that a biologically plausible decoder should be causal: only past neural activity can be taken into account, not future activity. Temporal averaging will, therefore, shift the population vector to reflect an estimate of the stimulus that lies in the past, unless the neurons anticipated future values of the stimulus. To make the population-vector estimate unbiased, the time window should be *T* = 2*τ*, where *τ* is the average ATI.

As shown in this study, the measured ATIs in the rat ADN are ideally suited to maximize the read-out accuracy. This is possible because the HD trajectory is sufficiently smooth to allow the extrapolation *θ*(*t* + *τ*) ≈ *θ*(*t*) + *ω*(*t*)*τ* so that HD cells can make a useful prediction of future HD values, reminiscent of predictive coding in sensory systems (Palmer et al., 2015). Random head movements with a frequency higher than 1/*τ* would make it impossible to significantly reduce the bias by anticipation. Indeed, the measured temporal correlations in the recorded HD signal are in the range of seconds and are thus much longer than the ATIs measured in the Papez circuit. The importance of anticipation is underlined by the fact that the bias contributes as much to the error in estimating the HD as neural variability itself, at least for model populations of HD cells with experimentally measured parameters (Figure 7).

Anticipatory firing might compensate for temporal delays that inevitably arise in sensory transduction (Berry et al., 1999) or as information is processed by a multi-layered system (Nijhawan and Wu, 2009); yet this explanation might belie anticipation’s true computational purpose. Compensating for delays does, for example, not explain why ATIs *increase* when rats are rotated passively (Bassett et al., 2003) – the delays across the Papez circuit should remain unaffected. Indeed, the increase in ATIs also runs counter to the hypothesis that motor-efference signals cause the ATIs; passive rotation eliminates the motor efference, which should cause the ATI to decrease, not increase. Yet a different experimental observation might have an important implication for decoding HD under passive rotation: firing rates in ADN decrease (Bassett et al., 2005). To counterbalance the resulting loss of information about HD in the ADN population, the read-out window *T* should increase (Burak and Fiete, 2012), and so should the ATI according to our framework. This hypothesis assumes that the ATI adapts to maintain an optimal read-out, which is, as yet, unproven.

If the anticipatory HD signal in the ADN were computed based on the read-out from another HD population, we expect even larger ATIs for that upstream population. As optimal ATIs change only slightly with system size (Figure 9A), ATI differences of around 25 *ms* are likely to occur along the hierarchy of the HD system. The staggered ATIs found in the Papez circuit – from *τ* = 40 − 75 *ms* in the rat lateral mammillary nuclei (LMN) to *τ* ≈ 25 *ms* in the ADN and retrosplenial cortex to zero lag in the postsubiculum (Taube 2007) – are consistent with this idea. Note that, in parallel, LMN activity could also be read out directly over a time window of 80 − 150 *ms* to yield a second estimate of current HD in addition to that provided by the ADN. Furthermore, the hypothesis of optimal population-vector decoding leads to testable predictions for the ATIs in other species, too, such as flies (Selig and Yayaraman, 2015; Kim et al., 2017) or bats (Finkelstein et al., 2015). All the information that is needed to calculate optimal ATIs for head direction can be derived from the movement statistics, single-neuron properties, and network size. If the movement statistics changes with the behavioral state, so should the ATIs. While cells in the dorsal subiculum of the bat change the encoding of head direction from pure to conjunctive coding when the animal switches from straight long-range navigation to rapid small-scale maneuvering (Finkelstein et al., 2018), the anticipatory behavior of these cells has not yet explicitly been addressed.

The HD system is a well-studied model, but it is but one example of how bias-variance trade-offs influence causal decoders of neuronal information. In many hierarchical processing streams in the nervous system, neurons must gather sufficient information to reliably perform computations while still adapting quickly enough to reflect changing inputs. Rather than being governed solely by biophysical limits on the speed of signal propagation, integration and action-potential generation, the time constants of dendritic and somatic integration as well as axonal propagation delays might also reflect decoding constraints. Further work is needed to elucidate how the requirement that the nervous system operate in real time shapes processing delays.

## Acknowledgements

This work was supported by the German Federal Ministry for Education and Research Grants 01GQ0440. We are grateful to G.Buzsáki for making data from Mizuseki et al. (2009) publicly available and thank A.Loebel and A.Mathis for stimulating discussions.

